# A murine *Wac* model exhibits phenotypes relevant to DeSanto-Shinawi Syndrome

**DOI:** 10.1101/2022.01.24.477600

**Authors:** April M Stafford, Maria Pacheco-Vergara, Katie L Uhl, Tara E Jager, Xiaopeng Li, Juhee Jeong, Daniel Vogt

## Abstract

Several monogenic syndromes are associated with neurodevelopmental changes that result in cognitive impairments, including autism, attention deficit hyperactivity disorder (ADHD) and seizures. Limited studies and resources are available to make meaningful headway into the underlying mechanisms that result in these symptoms. One such example, DeSanto-Shinawi Syndrome (DESSH), is a rare disorder caused by mutations in the *WAC* gene. Those diagnosed with DESSH experience craniofacial alterations as well as cognitive symptoms that include autism, ADHD and seizures. However, no thorough studies from a mammalian model exist to understand how these changes occur. To overcome this, we generated constitutive murine *Wac* mutants and assessed phenotypes that are relevant to humans diagnosed with DESSH. *Wac* mutants have craniofacial, anatomical, behavioral and seizure susceptibility that are relevant to DESSH; this new model is suited to study some of the core symptoms of DESSH and the biology of *Wac*.

## Introduction

Dysfunction of the WW domain-containing adaptor with coiled coil gene, *Wac*, underlies characteristic symptoms of Desanto-Shinawi syndrome (DESSH) [1]. DESSH was initially characterized by a combination of hypotonia, cranio-facial dysmorphic features, developmental delay and neurological/neuropsychiatric symptoms [1]. Neurological and neuropsychiatric symptoms include attention deficit hyperactivity disorder (ADHD), autism spectrum disorder (ASD) and seizures [1–5]. *Wac* is considered a high confidence ASD risk gene due to the elevated genetic variants discovered in ASD populations [6]. Despite these important details little is known about how *Wac* functions in the brain and whether a mammalian model could be generated to explore the causes of these associated symptoms.

The *Wac* gene encodes a protein that contains WW and coiled coil domains, motifs important for protein/protein interactions. Previous reports have demonstrated a multitude of cellular roles for the WAC protein, including positive regulation of mammalian target of rapamycin (MTOR), mitosis, transcriptional initiation and autophagy [7–10]. These unique roles are likely due to the distinct protein/protein interactions that WAC mediates in different tissues/cells by the WW and coiled coil domains. However, many of these studies have been performed in cell lines or models that are very different from a mammalian brain. Whether the same interactions or unique events occur in the mammalian brain is still unknown but will be important for investigating the role of WAC dysfunction and the symptoms associated with DESSH.

We generated a constitutive *Wac* deletion mouse model to assess whether loss of *Wac* could recapitulate the various symptoms associated with DESSH. While loss of both *Wac* alleles led to early embryonic lethality, we were able to generate and study mice with loss of one *Wac* allele, which is predicted to be analogous to most humans diagnosed with DESSH. *Wac* heterozygous (Het) mice exhibited dysmorphic craniofacial features, decreased expression of GABAergic interneuron proteins, susceptibility to seizures and behaviors implicating decreased learning and memory, suggesting some relevance to the underlying biology that causes symptoms of DESSH. Importantly, these data suggest that this mouse model may be a resource for future experiments to investigate therapeutics for the various symptoms of DESSH and could be used for further exploration of the biological mechanisms underlying this rare syndrome.

## Methods

### Animals

We thank the Wellcome Trust Sanger Institute Mouse Genetics Project (Sanger MGP) and its funders for providing the conditional mutant mouse gamete (C57BL/6N-Wac<tm2c(EUCOMM)Wtsi>/Wtsi). Funding information may be found at www.sanger.ac.uk/mouseportal, associated primary phenotypic information at www.mousephenotype.org and data about the development of these technologies [11–15]. Sperm harboring the *Wac* conditional allele was used to fertilize C57BL6/N donor eggs; progeny were then genotyped via polymerase chain reaction (PCR). We next bred *Beta actin-Cre* mice [16] with *Wac*^*Flox*^ mice to generate wild type (WT) and constitutive Het mice. After germline recombination, *Wac* Het mice were backcrossed and bred with CD-1 mice for at least three generations before being used for experiments, which were performed under the approval of Michigan State University’s Campus Animal Resources.

### Behavior

Paradigms tested ambulatory and working memory behaviors that were previously optimized [17, 18]. 6-8 week aged mice of both sexes were tested; no differences were observed in any assay between sexes and data represent both groups. The persons performing the behaviors and analyses were blinded to the genotypes.

#### Open field

Individual mice were placed in an open chamber with a center region designated by a 10 × 10 cm^2^ area and the remaining area to the wall designated as periphery. Mice roamed freely for ten minutes while being tracked by an overhead camera. The total distance travelled, and the mean speed, were analyzed using ANY-maze software as well as time spent in center versus periphery.

#### Y-maze

Individual mice were placed at the end of one arm of a Y-maze and allowed to explore for five minutes while being filmed by an overhead camera. Entries into all arms were noted (four paws need to be inside the arm for a valid entry) and a spontaneous alternation is counted if an animal enters three different arms consecutively. % of spontaneous alternation were calculated according to following formula: [(number of alternations) / (total number of arm entries − 2)] × 100.

### Cranio-facial analyses

The skulls were stained with Alizarin red for bone as previously published [19]. Quantification of the suture and fontanel areas and the skull width was performed from photographs using ImageJ.

### Genotyping

Primers to detect the recombined allele of the *Wac* genetic locus: Forward 5’-AGCTATGCGTGCTGTTGGG-3’ and Reverse 5’-CAAATCCCACAGTCCAATGC-3’. Thermocycling conditions were: 95ºC 3 minutes, (95ºC 30 seconds, 58ºC 30 seconds, and 72ºC 45 seconds for 35 cycles), 72ºC 3 minutes. Sanger sequencing to validate the recombined locus was performed by GeneWiz using the same primers on gel-purified PCR products.

### Immuno-fluorescent staining

At P30, mice were transcardially perfused with phosphate-buffered saline (PBS), followed by 4% paraformaldehyde. The brains were then removed and postfixed in PFA for 30 minutes. Brains were transferred to 30% sucrose for cryoprotection overnight after fixation and then embedded in optimal cutting temperature compound before coronally sectioned at 25μm via cryostat. Sections were permeabilized in a wash of PBS with 0.3% Triton-X100, then blocked with the same solution containing 5% bovine serum albumin. Primary antibodies were either applied for 1 hour at room temperature or overnight at 4ºC, followed by 3 washes. Secondary antibodies were applied for 1-2 hours at room temperature, followed by 3 washes and cover slipped with Vectashield. Primary antibodies included: goat anti-IBA1 (1:500, NB100-1028 Novus Biologicals), mouse anti-LHX6 (1:200, sc-271433 Santa Cruz Biotechnology), mouse anti-NeuN (1:400, MAB377 MilliporeSigma), rabbit anti-OLIG2 (1:400, AB9610 MilliporeSigma), rabbit anti-PROX1 (1:500, ab199359 Abcam), rabbit anti-PV (1:400, PV27 Swant), rabbit anti-S100beta (1:500, 15146-1-AP Proteintech), rat anti-SST (1:200, MAB354 MilliporeSigma). Secondary Alexa-conjugated fluorescent secondary antibodies (488 and 594 wavelengths) were from Thermo-Fisher and used at a 1:300 dilution.

### Microscopy

Fluorescent images were acquired using a Leica DM2000 microscope with a mounted monochrome DFC3000G camera. Fluorescent images were adjusted for brightness/contrast and merged using Image-J software. The skulls were photographed with Nikon SMZ1500 stereomicroscope and Nikon DSRi1 camera.

### PTZ induced seizure assessments

Pentylenetetrazole (PTZ) was prepared fresh for each experiment. 1 gram of PTZ powder was dissolved into 250 ml of PBS. PTZ solution was injected intraperitoneally at a dose of 50 mg of PTZ per gram of mouse body weight. This dose was determined to be subthreshold for inducing a seizure in most WT mice, but resulting in motor twitches and freezing behavior. After injection, mice were placed in a clean cage and monitored for 20 minutes. Seizure severity was scored based on a modified Racine scale [20], which followed these criteria: 1) Freezing or idle behavior; 2) Muscle twitches and/or rhythmic head nodding; 3) Tail arching over the mouse’s backside accompanied by the mouse assuming a hunched posture; 4) Forelimb clonus; 5) Generalized motor convulsions accompanied by loss of balance (resembles tonic-clonic seizure) but with a recovery to idle or normal behavior and not advancing to level 6/7 phenotypes; 6) Uncontrolled running, jumping and hyperactivity; 7) Full body extension of all limbs; 8) Death.

### Western blots

P30 somatosensory cortices were dissected and frozen on dry ice. Next, they were lysed in RIPA buffer containing protease and phosphatase inhibitors and combined with Laemmli buffer containing 2-Mercaptoethanol and incubated at 95ºC for 5 minutes to denature the proteins. Equal amounts of protein lysates were separated on 10% SDS-PAGE gels and then transferred to nitrocellulose membranes. The membranes were washed in Tris-buffered saline with Tween-20 (TBST) and then blocked for 1 hour in TBST containing 5% non-fat dry milk (blotto, sc-2324 SantaCruz biotechnology). Membranes were then incubated with primary antibodies overnight at 4ºC, washed 3 times with TBST, incubated with secondary antibodies for 1 hour at room temperature and then washed 3 more times with TBST. Membranes were next incubated in ECL solution (BioRad Clarity substrate 1705061) for 5 minutes and chemiluminescent images obtained using a BioRad Chemidoc™ MP imaging system. Antibodies included: rabbit anti-WAC (1:2000, ab109486 Abcam), rabbit anti-pS6, rabbit anti-S6 and rabbit anti-GAPDH (1:4000, 5364, 2217 and 2118 Cell Signaling Technology), goat anti-rabbit HRP (1:4000, 170-6515 BioRad).

### Quantification and statistical analysis

Statistical analyses were performed using Prism version 6, a p value of < 0.05 was considered significant. For parametric measures of two groups, a two-tailed T-test was performed.

## Results

### Validation of a constitutive murine *Wac* mutant model

We first obtained sperm from the International Mouse Phenotyping Consortium (IMPC), in which the *Wac* murine locus on chromosome 18 was genetically modified to include flanking *loxP* (Flox) sites of exon 5 (Schema, Figure 1a). *Wac* flox founders were generated at the Michigan State University Genomics Core and validated via PCR genotyping. To produce experimental progeny, *Wac* flox mice were crossed to *beta-actin-Cre* mice, who express Cre in germ cells [16]. We were able to produce both wild type (WT) and constitutive heterozygous (Het) progeny (genotyping PCR shown in Figure 1b) but never any live knockouts when Het mice were bred together. Finding were consistent with previous data from the IMPC that constitutive *Wac* knockouts are embryonic lethal.

**Figure 1:**
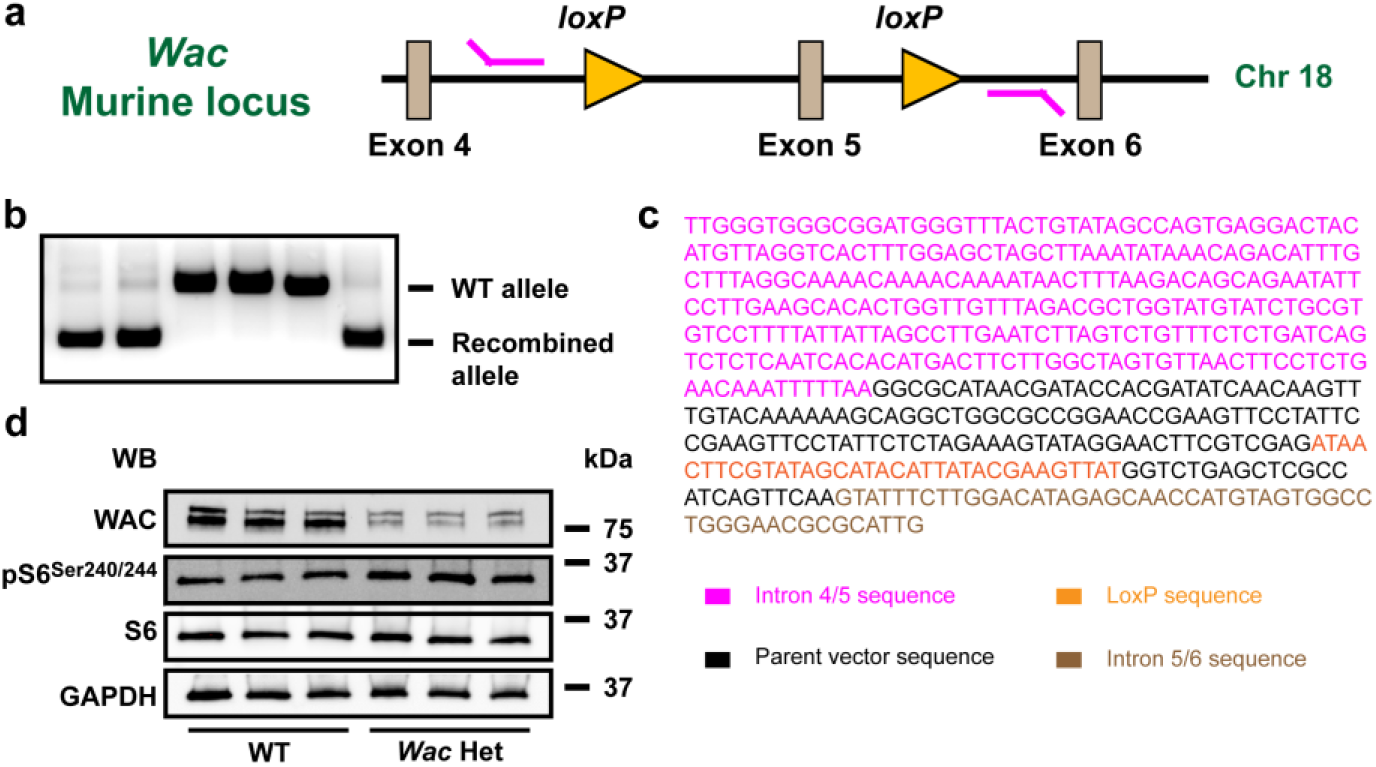
*Wac* genetic locus and validation. The floxed mouse *Wac* genetic locus and genotyping primers are shown in (a). Genotyping primers (magenta lines) reside external to the *loxP* sites (orange triangles) that flank exon 5 and portions of surrounding introns. (b) DNA gel showing genotyping results for WT and recombined (Het) *Wac* alleles. (c) The recombined DNA band was Sanger-sequenced and found to contain a single recombined *loxP* site (orange) flanked by intronic DNA sequences (magenta and brown text); additional text matches the parent DNA vector used to generate the locus. (d) Western blots of P30 somatosensory cortices to detect WAC, the MTOR target, pS6, total S6 and GAPDH proteins in 3 pairs of P30 brains. Abbreviations: (Chr) chromosome, (WB) western blot and (kDa) kilodaltons.

To validate that our genotyping results were accurately detecting the *Wac* recombined locus in the mutant animals, we performed Sanger sequencing on the DNA PCR product presumed to represent the recombined locus in Het genotyping samples. Indeed, sequencing indicated that DNA from the surrounding introns and a single *loxP* site corresponded to the region on chromosome 18 within the *Wac* locus but lacked a second *loxP* site, exon 5 and flanking intronic sequences between the *loxP* sites (Figure 1c). Finally, we assessed postnatal day (P)30 cortical brain tissue for WAC protein in three sets of WTs and Hets to determine if the protein was decreased in the mutant; we observed WAC protein loss in all the Hets (Figure 1d). Interestingly, we did not detect changes in *Wac* transcript, suggesting possible compensation at the WT locus or the presence of an alternatively spliced isoform (data not shown). Finally, we probed for the MTOR activation marker, phosphorylated ribosomal subunit S6, from the same protein lysates that had depleted WAC. In contrast to WAC’s role in drosophila [9], we did not find a role for mammalian WAC in promoting MTOR activation (Figure 1d), although a non-significant trend in increased activation was noted. These data suggest unique mechanisms may underlie how mammalian WAC results in DESSH symptoms.

### Craniofacial changes in neonatal and adult *Wac* Het mice

Since craniofacial dysmorphism is commonly observed in those diagnosed with DESSH [1–3, 5, 21–25], we wanted to understand if a reduction in *Wac* dosage would recapitulate some of these observations in mice. To this end, postnatal day (P)0 and P30 mouse skulls were examined. During neonatal ages, a widening of the fontanels and sutures along the midline of the calvaria was noted (Figure 2a, 2b). While both the anterior and posterior regions of the calvaria were affected, the anterior region had the greater change (Figure 2a, 2b, 2e, 2f with pseudo-colored areas in 2a’, 2b’; anterior p = 0.0001, posterior p = 0.014). By P30, there remained a noticeable gap in the interfrontal suture of *Wac* Het. Concomitantly, the width of the skull across the frontal bones was significantly increased in *Wac* Hets, while the width across the parietal bones was not altered (Figure 2c, 2d, 2g, 2h, p = 0.0022 Frontal width). These data reveal that this mouse model has craniofacial changes that may recapitulate some features observed in DESSH.

**Figure 2:**
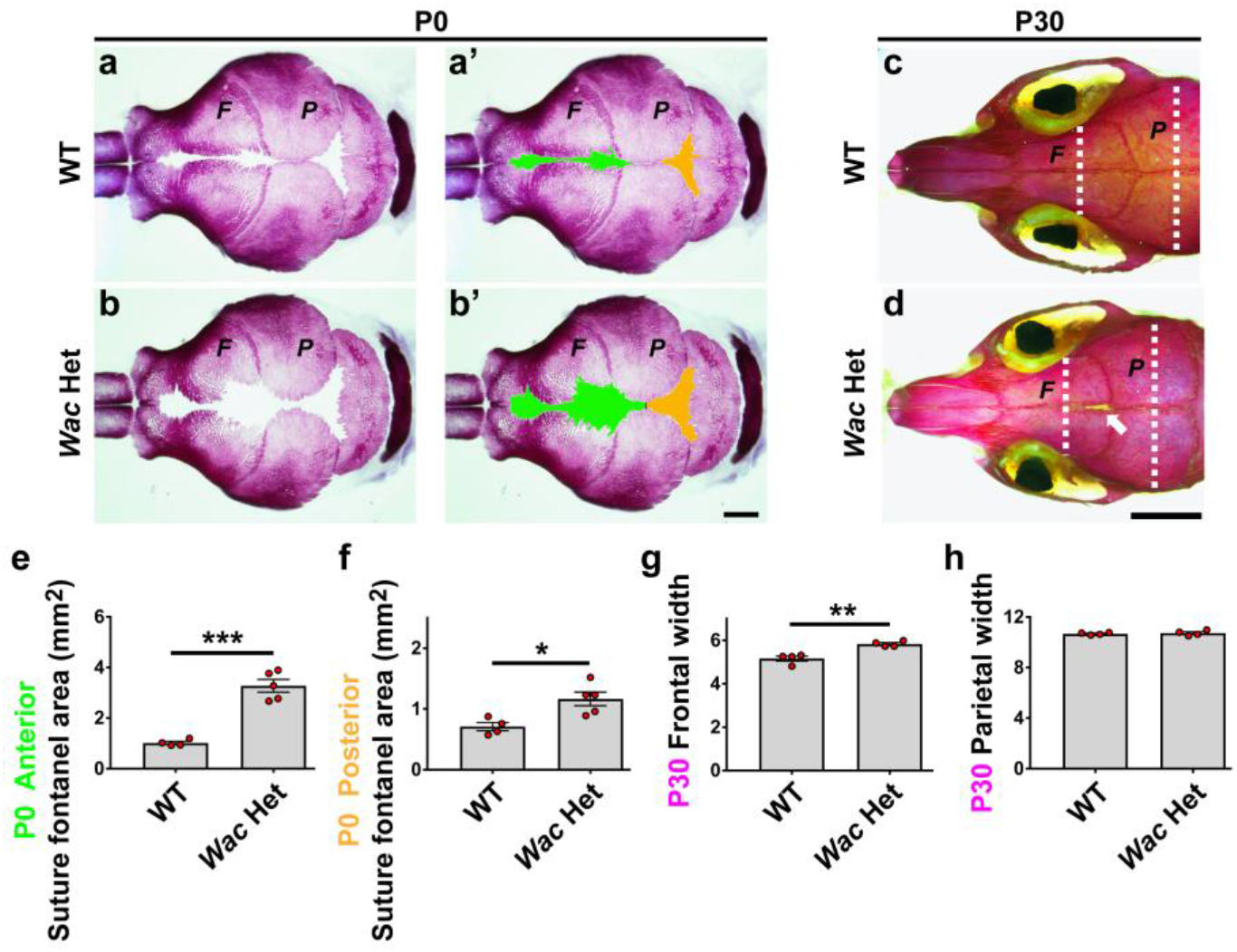
Craniofacial changes in *Wac* Het mice. (a, b) Dorsal views of the P0 calvaria stained with Alizarin red for bone. (a’, b’) The suture and fontanel areas in (a) and (b) are pseudo-colored green (anterior) and orange (posterior). (c, d) Dorsal views of P30 WT and *Wac* Het skull bones; arrow (d) points to a gap in the interfrontal suture. (*F*) Frontal bone and (*P*) Parietal bone; white dashed lines are widths measured in (g, h). (e, f) Quantification of P0 fontanel suture areas pseudo colored in a’ and b’. Quantification of the skull width across the frontal (g) and parietal (h) bones at P30 in WTs and *Wac* Hets. Data are expressed as the mean ± SEM, P0 n = 4 (WT), n = 5 (Het); P30 n = 4 (WT), n = 4 (Het) biological replicates. * p<0.05, ** p<0.01 and *** p<0.001. Scale bars: (b’) = 1mm, (d) = 4mm.

### Subtle behavioral changes in *Wac* mutant mice

Whether loss of *Wac* may impact cognitive behavior was also a goal as those diagnosed with DESSH exhibit some intellectual disability [3, 5, 22, 23, 25] and a drosophila loss of function model has shown deficits in learning [23]. A panel of behavioral tests were used to assess movement as well as working memory. We first used the open field test to assess ambulatory and potential anxiety-like behaviors (Schema Figure 3a). WT and Het mice performed similarly and no signs of anxiety or impaired movement were observed, or in preference for the central or peripheral domains (Figure 3b)

**Figure 3:**
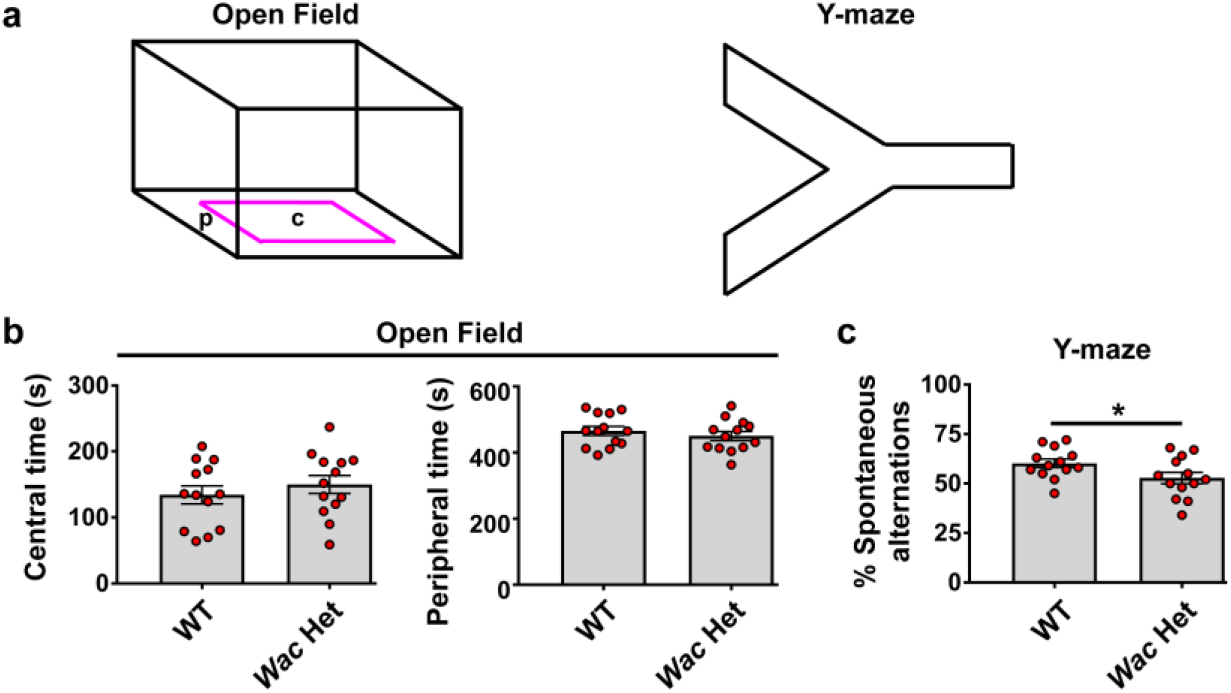
*Wac* hets exhibit reduced working memory. (a) 6-8 week aged WT and *Wac* Het mice were tested in the Open Field and Y-maze paradigms (a); magenta square delineates the central, **c**, from peripheral, **p**, regions measured. Mice were assessed for their time spent in the center or periphery in the open field (b), with no differences noted. In the Y-maze, *Wac* hets had decreased spontaneous alternations (c). Data are expressed as the mean ± SEM; n = 13, all groups; * p<0.05.

To assess working memory, we employed the Y-maze test [17, 26] to measure the ability of WT and *Wac* Het mice to spontaneously alternate into novel arms. This behavior takes advantage of the mouse’s novelty seeking while working to remember the most recent arms explored. As expected, WT mice exhibited a high rate of spontaneous alternations, i.e. selecting a novel arm during exploration of the maze, the *Wac* Het mice had a significantly lower rate (Figure 3c, p = 0.0474), suggesting a potential impairment in working memory.

### Normal numbers of major brain cell types

To determine if any major cell types of the brain were altered by loss of *Wac*, we probed for different classes of neurons, as well as oligodendrocytes, astrocytes and microglia in the somatosensory cortex (Figure 4a). The total number of neurons were assessed by labeling for NeuN, while oligodendrocytes, astrocytes and microglia were labeled with OLIG2, S100beta and IBA1, respectively. No differences in these cell types were observed between the genotypes (Figure 4a). There were also no gross changes in cell distribution that we could detect in other brain areas (data not shown).

**Figure 4:**
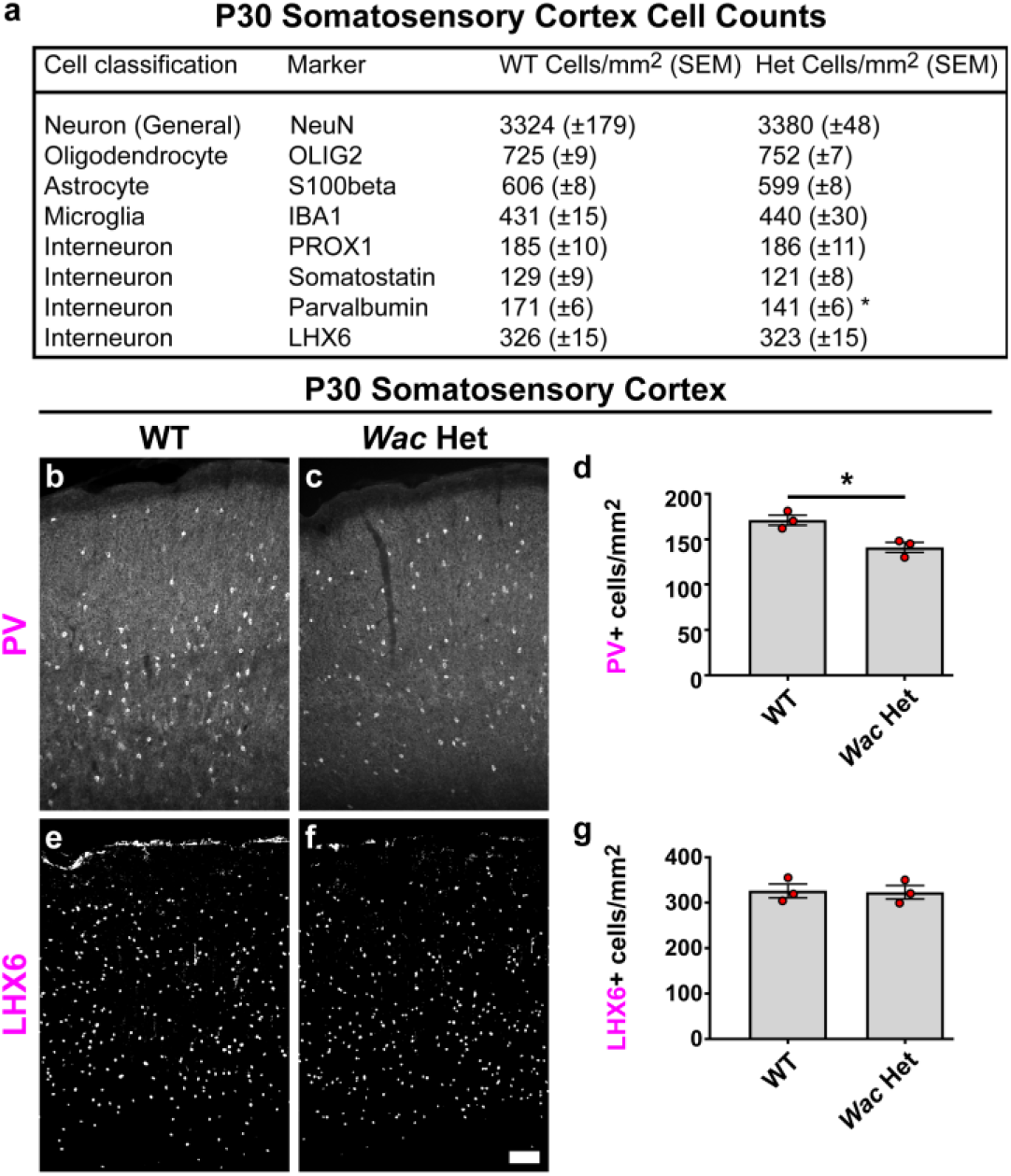
Loss of *Wac* results in a decrease in PV expression. (a) P30 cell density counts from the somatosensory cortices of WT and *Wac* Het mice; most cell types are not changed. (b, c) immunofluorescent images of PV and cell density quantification (d), showing a decrease in expression. (e, f) immunofluorescent images of LHX6 and cell density quantification (g), showing no change. Data are expressed as the mean ± SEM; n = 3 biological replicates, all groups. * p<0.05. Scale bar in (f) = 100μm.

### Distinct changes to GABAergic cell types in *Wac* mutant mice

Although there were no gross changes in total neuron numbers, some neuron cell types are sparse and may not be well represented by this approach. These include GABAergic cortical interneurons (CINs), a diverse group of inhibitory neurons whose dysfunction has been implicated in other monogenic syndromes [27–32]. To assess whether loss of *Wac* may impact CINs, we stained for 3 mostly non-overlapping groups of CINs, those expressing parvalbumin (PV), somatostatin (SST) and prospero homeobox 1 (PROX1). While the cell density of SST and PROX1 CINs were not different (Figure 4a), there was ∼18% decrease in CINs expressing PV in the Het (Figure 4a-4d, p = 0.019). Thus, while the numbers of core brain cell types are normal, PV expression is uniquely impacted by decreased *Wac* levels. Finally, to probe whether the loss of PV expression may be due to loss of cells or simply in the expression of PV, we assessed for the MGE-lineage CIN marker, LHX6, of which ∼half are PV+ in the somatosensory cortex. We found no decrease in LHX6 cell numbers (Figure 4e-4g), suggesting the loss of PV is not likely due to cell loss but rather PV expression; a similar phenotype was reported in another syndromic mouse model (Vogt et al., 2018).

### Elevated seizure susceptibility in *Wac* heterozygous mice

Since PV expression was decreased and seizures have been observed in some individuals diagnosed with DESSH [1, 3, 4, 23, 25], we wanted to determine if constitutive loss of one *Wac* allele was sufficient to increase seizure susceptibility. We did not observe any spontaneous seizures by P30. However, to further assess, the GABAA receptor antagonist, pentylenetetrazol (PTZ), was administered intraperitoneal at a dose (50mg PTZ per kilogram of mouse body weight); this subthreshold seizure dose caused freezing and mild muscle twitching behavior in WT mice but rarely elicited an observed seizure. We hypothesized that if loss of *Wac* promoted a shift favoring brain excitation/inhibition that *Wac* hets would exhibit more severe seizure behaviors by P30.

PTZ was administered and mice were observed for 20 minutes (Schema, Figure 5a); most mice fully recovered by 15 minutes and all WT mice did not exhibit any behavioral changes by twenty minutes post injection. We also separated females and males due to subtle elevated responses in male WT mice administered PTZ compared to females. PTZ led to subtle behavioral changes in both female and male WT mice but rarely elicited a seizure (Figure 5b and 5c). However, both sexes of *Wac* hets exhibited elevated seizure behaviors, including hunched body posture, arching tail and forelimb clonus as well as succumbing to seizures (Figure 5b and 5c; females p = 0.0008 and males p = 0.0047). Thus, loss of *Wac* leads to increased seizure susceptibility in mice, consistent with one of the comorbidities observed in some individuals diagnosed with DESSH.

**Figure 5:**
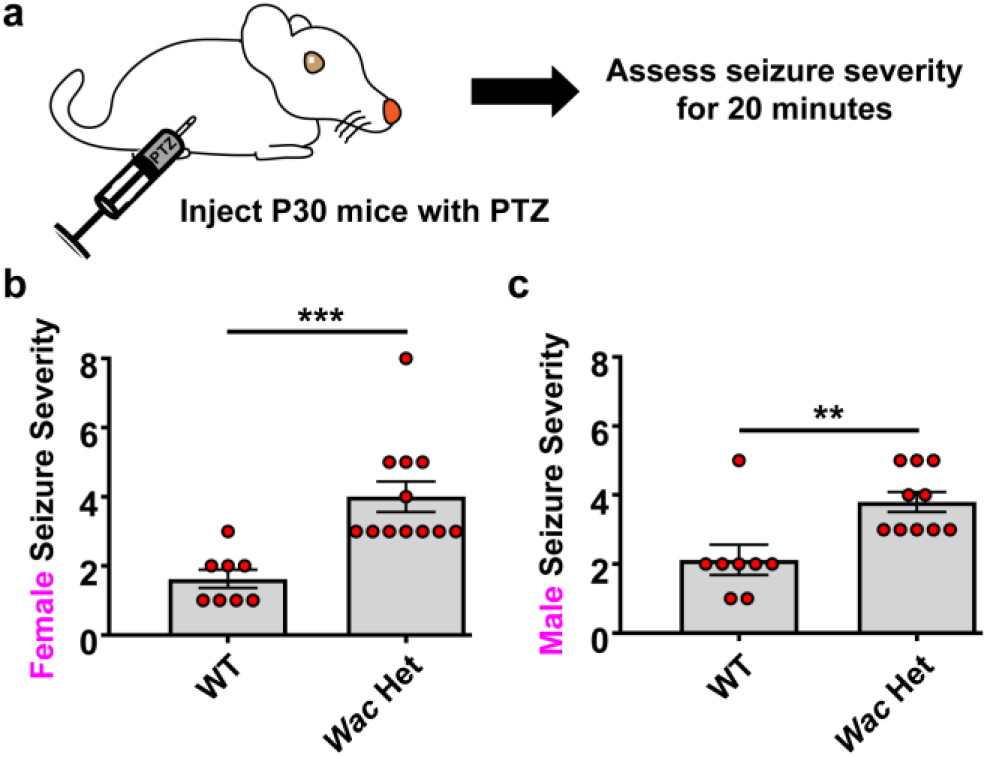
*Wac* depletion leads to elevated seizure susceptibility. (a) Schema depicting the seizure induction by PTZ. Briefly, P30 mice were administered PTZ intraperitoneally and then assessed for the highest seizure severity score over the course of 20 minutes. The maximum seizure severity score over 20 minutes in females (b) and males (c) demonstrate an increased sensitivity to seizures in the *Wac* Hets. Data are expressed as the mean ± SEM; for females, n = 8 (WT), n = 12 (Het) and for males, n = 8 (WT), n = 10 (Het) biological replicates. ** p<0.01 and *** p<0.001.

## Discussion

DESSH is a monogenic syndrome that results in developmental delay, craniofacial dysmorphic features and neurological changes that may underlie the elevated co-diagnoses of ASD and ADHD. Previous work implied that dysfunction of the *Wac* gene is associated with DESSH symptoms, thus we sought to test if genetic depletion of the *Wac* gene could recapitulate some of the reported symptoms and provide a novel mammalian model to study DESSH. To this end, we deleted *Wac* using a germline expressing Cre to test the impact of loss of function at the heterozygous level. Overall, these data provide evidence that this *Wac* mouse model does indeed show evidence of relevant phenotypes to humans diagnosed with DESSH and could prove to be a great tool in understanding the biological mechanisms underlying the syndrome as well as testing future therapeutics and interventions to remedy the symptoms of DESSH. While these data show key phenotypes that are of relevance to DESSH they have not covered all the symptoms associated with the syndrome, which will be a major task in the future.

A common symptom in DESSH are dysmorphic facial features REFs. Upon examination of craniofacial development in *Wac* Het mice, there was a broadening of the fontanels and sutures along the midline of the calvaria at birth, with a greater difference in the anterior region. By P30, only the frontal bone region in the anterior part of the calvaria exhibited increased width in the Hets, while the posterior, parietal bone region was not different. These data suggest that frontal cranial development is impacted greater and since posterior regions are normal by P30, these changes are not likely due to a general increase in head growth. This is important because those diagnosed with DESSH have dysmorphic facial changes, including a widening of the forehead, which suggests this mouse model has craniofacial features relevant to DESSH.

Intellectual disability is also found in those diagnosed with DESSH. One feature of this may include difficulty in some learning and memory tasks that can be modeled in mice (herein) and other model organisms [23]. We found loss of *Wac* resulted in behavioral deficits that measure short term memory. These data imply that this murine model could be used in future studies to assess therapeutic efficacy of treatments on memory. Moreover, other, more specific tests, may be helpful as well as using this model to understand the high rate of ADHD in the DESSH population and other comorbidities.

Molecular changes in key brain markers were probed that may be key to discovering mechanisms underlying the symptoms associated with DESSH. Overall, brain cell types and their numbers were not grossly impacted by loss of *Wac*. This is important because it means that irreversible cell changes have not occurred in the populations we assessed and their manipulation with future therapeutics could be promising if more subtle phenotypes are discovered in the future. However, we did find that one population of inhibitory interneurons was impacted in *Wac* Hets, those expressing PV. While subtle, these changes may be important for the behavioral changes and the increased susceptibility to seizures that we also found to occur in these mice. We and others have observed subtle decreases of PV expression in other syndromic mouse models, including *Cntnap2* (Peñagarikano et al., 2011; Vogt et al., 2018; Paterno et al., 2021), suggesting this may be a common feature in some monogenic syndromes.

Finally, we asked whether loss of *Wac* led to a susceptibility to seizures, as some diagnosed with DESSH have seizures/epilepsy [1, 3, 4, 23]. Indeed, we found that a subthreshold doze of PTZ to inhibit GABAA receptors caused *Wac* Het mice to seize when their WT littermates often did not. While we did not notice spontaneous seizures in the Hets, their reaction to PTZ suggest that their brains are shifted towards being more excitable. It is also possible that this more excitable brain may influence the expression of PV, as we and others have found that brain activity can alter PV expression [31, 35]. Thus, it is still unclear whether the loss of PV leads to seizures in our model or if the elevated brain excitability could lead to changes in PV expression. No matter the cause, others have noted the importance of PV expression on behaviors relevant to ASD [36], suggesting that approaches to restore PV expression may be a future therapeutic to alleviate some DESSH symptoms. In total, we have validated a new mouse model that has features relevant to some of the symptomatology of DESSH. These phenotypes are a first attempt to understanding this complex rare disorder and offer a tool to the field to probe novel mechanisms and therapeutics that could benefit those diagnosed.

## Acknowledgements

**AMS, DV, KU, TEJ** and **XL** were funded by the Michigan State University/Spectrum Health Corporation; **MP-V** and **JJ** were funded by R01 DE026798; **KU, TEJ** and **XL** were also funded by the Cystic Fibrosis Foundation LI19XX0, the Cystic Fibrosis Research Inc., R01 HL153165-01A1.

## Author contributions

**AMS** and **DV** validated the mouse strain, processed tissue for staining and performed PTZ studies. **MP-V** and **JJ** performed craniofacial experiments. **KU, TEJ** and **XL** performed *Wac* expression analyses. **DV** did western blots. **AMS** performed behavior experiments. All authors edited and approved the manuscript.

